# Sphingolipid Biosynthesis Inhibition As A Host Strategy Against Diverse Pathogens

**DOI:** 10.1101/2020.04.10.035683

**Authors:** Sandeep Kumar, Jinmei Li, Jiwoong Park, Sydney K. Hart, Niki J. Song, Damon T. Burrow, Nicholas L. Bean, Nicholas C. Jacobs, Ariella Coler-Reilly, Anastasiia Onyshchenko Pendergrass, Tanya H. Pierre, India C. Bradley, Jan E. Carette, Malini Varadarajan, Thijn R. Brummelkamp, Roland Dolle, Tim R. Peterson

## Abstract

Chloroquine is an anti-malarial and immunosuppressant drug that has cationic amphipathic chemical properties. We performed genome-wide screens in human cells with chloroquine and several other widely used cationic amphipathic drugs (CADs) including the anti-depressants, sertraline (Zoloft) and fluoxetine (Prozac), the analgesic nortriptyline (Pamelor), the anti-arrhythmic amiodarone (Cordarone), and the anti-hypertensive verapamil (Calan) to characterize their molecular similarities and differences. Despite CADs having different disease indications but consistent with them sharing key chemical properties, we found CADs to have remarkably similar phenotypic profiles compared with non-CADs we and others have previously screened (1–5). The most significant genetic interaction for all CADs was the initiating step in sphingolipid biosynthesis catalyzed by serine palmitoyltransferase (SPT). A comparison of genome-wide screens performed with diverse pathogens from viruses, bacteria, plants, and parasites including Ebola (6), adeno-associated virus AAV2 (7), HIV (8), Rotavirus (9), Influenza A (10), Zika virus (11), Picornavirus (12), Exotoxin A (13), Cholera toxin (14), Type III secretion system and Shiga toxin (15, 16), Ricin toxin (17), and Toxoplasma gondii (18) showed SPT as a top common host factor and 80% overlap overall in top hits specifically with CADs. Potential sphingolipid-mediated mechanisms for the host response- and virulence-modulating effects of CADs involve autophagy and SERPINE1/PAI-1 (plasminogen activator inhibitor-1). Chloroquine has recently shown potential as an anti-viral agent for the novel coronavirus SARS-CoV-2, the causative agent of COVID-19 respiratory disease (19, 20). Our study demonstrates that numerous readily available drugs molecularly function highly similar to chloroquine, which suggests they might be considered for further pre-clinical investigation in the context of SARS-CoV-2. More generally, our work suggests the diverse pathogen mitigating potential of drugs that inhibit host sphingolipid biosynthesis such as CADs.

**Brief Summary:** Our study demonstrates that numerous readily available drugs molecularly function highly similar to chloroquine, which suggests they might be considered for further pre-clinical investigation in the context of SARS-CoV-2.

## MAIN

Chloroquine is a cationic amphipathic drug (CAD). Many other United States Food and Drug Administration (FDA) approved drugs such as anti-depressants, sertraline (Zoloft), fluoxetine (Prozac), the analgesic nortriptyline (Pamelor), the anti-arrhythmic amiodarone (Cordarone), anti-hypertensive verapamil (Calan), and the anti-estrogen tamoxifen (Novladex) also have these chemical properties. Small molecules that have these properties are also referred to as lysosomotropic because protonation of their ionizable amines traps them in lysosomes (21). CADs are known to cause phospholipidosis (increased intracellular phospholipids) and to induce autophagy – a process of cell catabolism (22, 23), but those are generally believed to be off-target effects given the micromolar concentrations used in vitro to elicit them. This might need further investigation, however, due to recent in vivo evidence showing that fluoxetine and amitriptyline (nortriptyline is its metabolite) induce autophagy at the concentrations at which they have antidepressant effects (24, 25).

Chloroquine has recently shown potential as an anti-viral agent for the novel coronavirus SARS-CoV-2, the causative agent of COVID-19 respiratory disease (19, 20). That chloroquine might be effective in protecting humans against both viruses and parasites suggested to us that chloroquine might act on the same host cellular machinery in both cases.

### Evidence that the initiating enzyme in sphingolipid biosynthesis serine palmitoyltransferase is central to the mechanism of action for diverse CADs

CADs can be distinguished from non-CADs based on having both high pKa’s and cLogP’s (Figure 1A) (26). cLogP refers to a compound’s partitioning in octanol vs. water, reflecting its hydrophobicity/hydrophilicity. The larger the value the more hydrophobic, the smaller the more hydrophilic. For example, CADs such as chloroquine have a cLogP ~4-5 and are relatively hydrophobic. This means they are more likely to cross the blood brain barrier and indeed many neuropsychiatric drugs are designed to have high cLogP’s to increase their central nervous system penetration. Negative cLogP values such as with the bisphosphonate anti-osteoporotic drug alendronate indicate the compound is hydrophilic. pKa refers to an acid dissociation constant. A value greater than 7 means the compound is basic, a value less than 7 means it is acidic. CADs have pKa’s ~9-10 and are weak bases.

**Figure 1:**
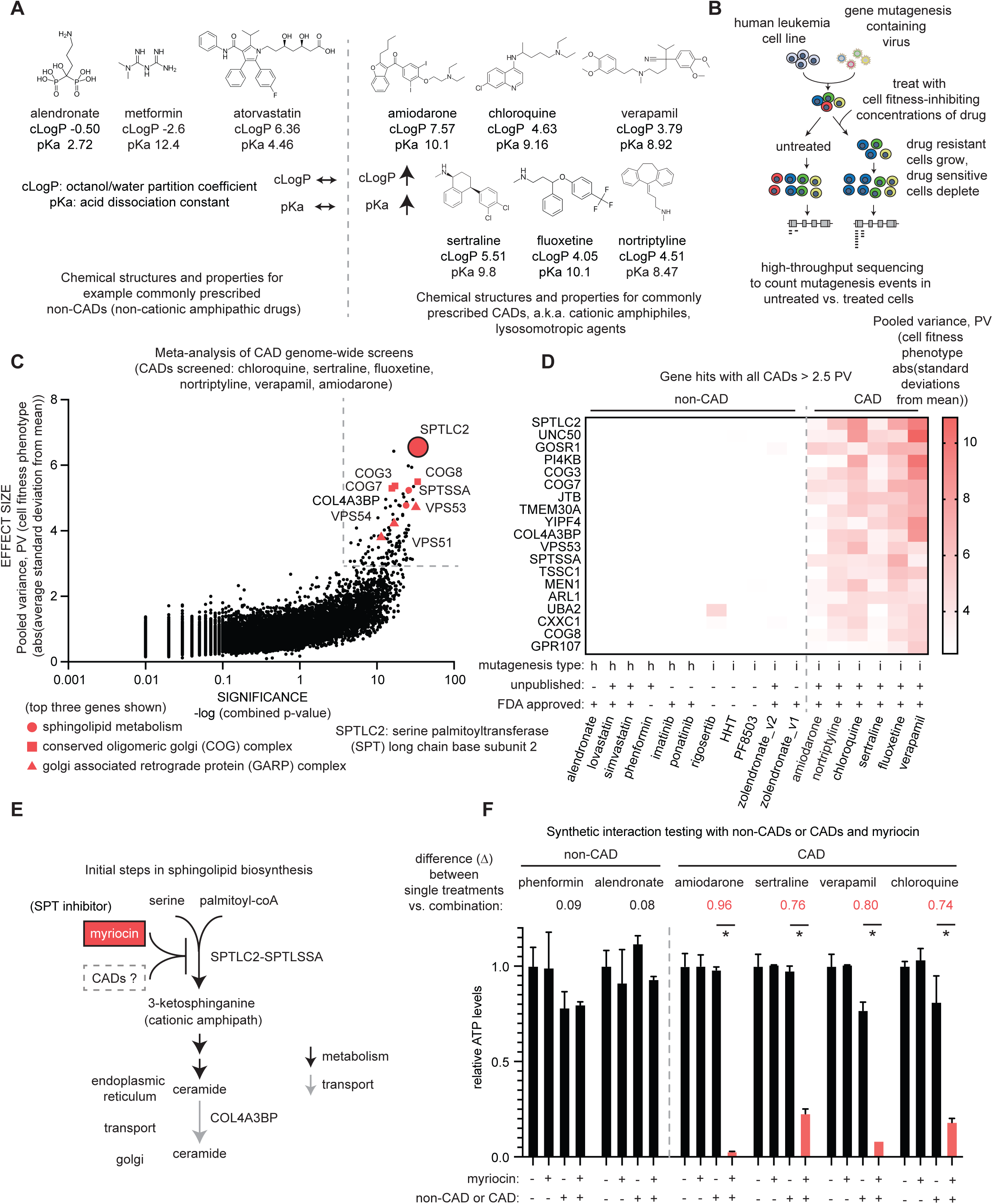
Genome-wide screens in human cells reveals inhibition of serine palmitoyltransferase as a potential mechanism of action for diverse CADs. **(a)** Chemical structures and properties of several FDA-approved cationic amphipathic drugs (CADs) and non-CADs. The following CADs: sertraline, nortriptyline, fluoxetine, verapamil, amiodarone, chloroquine and non-CADs: alendronate, metformin, atorvastatin are shown. CADs have relatively high partition coefficients (cLogP) and acid dissociation constants (pKa). A cLogP of ~4-5 is relatively hydrophobic. A pKa value of ~9-10 is weakly basic. **(b)** Schematic of the genome-wide screening process. A mutagenesis approach using CRISPR interference (CRISPRi) was used for CADs at GI50 concentrations of drug. GI50 refers to the concentration of drug that inhibits cell fitness by 50% relative to untreated cells. **(c)** Meta-analysis of CAD screens. Mutagenesis type “h” refers to haploid gene-trap (77). Mutagenesis type “i” refers to CRISPRi (14). Zolendronate_v1 and v2 refer to biological replicates. V1 was previously published (2). Pooled variance (PV) is the absolute value of the aggregated phenotype strength/effect size for a gene knockdown with all CADs that were tested. P-values for individual screens were ‘combined’ by FDR-adjusting for multiple comparisons. 18,191 human genes were analyzed, though 4,187 genes were not plotted because they have 0 values and the log of 0 doesn’t exist. GI50 drug concentrations of CADs used in CRISPRi screens were as follows: fluoxetine 30 µM, nortriptyline 30 µM, chloroquine 100 µM, verapamil 250 µM, amiodarone 50 µM, sertraline 9µM. Individual screen results are provided in Table S1. **(d)** Scoring of CAD top hits in each CAD and non-CAD screen. Comparisons of the phenotype scores for the 19 genes that had PV > 2.5 for all six CADs we tested vs. their PV’s with 10 non-CADs (alendronate, lovastatin, simvastatin, phenformin, HHT, PF8503, imatinib, ponatinib, zoledronate). Dotted lines reflect thresholds of PV > 2.5 and a FDR-adjusted p-value < 3.3E-4 for all six screens having individual p-values < 0.05. (**e**) Initial steps in the sphingolipid biosynthesis pathway and the design of a synthetic interaction experiment involving a combination of myriocin and a CAD. Serine palmitoyltransferase is the known site of action for myriocin and is our hypothesized site of action for CADs. **(f)** Synthetic interaction testing between SPT and CADs. The synthetic interaction strength between SPT and CADs was tested with 100nM myriocin, which inhibits all three SPTs (SPTLC1, SPTLC2, SPTLC3). Non-CAD and CAD doses used minimally inhibit cell fitness on their own based on data in Figure S1a. Doses used are as follows: non-CADs – 100µM alendronate, 50µM phenformin; CADs – amiodarone 15µM, verapamil 60µM, chloroquine 20µM, sertraline 11µM. * P-value < 0.05, two-way ANOVA for all controls vs. experimental samples comparing single agents (myriocin or CAD) vs. their combination (myriocin + CAD). Detailed information on statistical tests and exact p-values for each comparison are provided in Table S1.

To molecularly compare several clinically important CADs including chloroquine and other FDA approved CADs (sertraline, fluoxetine, nortriptyline, amiodarone, verapamil), we performed genome-wide screens in a human K562 chronic myelogenous leukemia cell line at concentrations that inhibit their growth 50% compared to untreated (GI50, <100 micromolar (µM) range, Figure S1A) using cell fitness as a phenotypic readout (Figure 1B). While our initial validation of sertraline hits suggested that we could detect strong genetic interactions in genome-wide screens with CADs (Figure S1B), there are valid concerns about whether using micromolar (µM) doses in cancer cell lines with cell fitness as a readout can give insight into in vivo biology for drugs such as these CADs, which are not indicated for cancer. However, we have previous demonstrated the success of this approach with the bisphosphonates (1, 2, 27), which suggest that it might yield meaningful results for the CADs as well (14).

A comparison of the six CADs we screened vs. genome-scale screens (as opposed to targeted screens, e.g., involving the kinome) we and others performed with non-CADs shows the specificity of the CADs signature (Figure 1C) (1–5). There were nineteen genes that scored on average an absolute value of at least 2.5 standard deviations away from the mean for the six CADs screens (hereafter referred to as pooled variance, PV) (Figure 1C, **Table S1**). Said another way, PV is the aggregated phenotype strength/effect size for a gene knockdown with all CADs we tested. The vast majority of top hits were endoplasmic reticulum (ER) and golgi localized factors including those that regulate sphingolipid metabolism and subunits of the conserved oligomeric golgi (COG) - COG3, COG7, and COG8 and golgi-associated retrograde protein (GARP) - VPS53, VPS54, VPS51 complexes (28, 29). The initiating enzyme in sphingolipid biosynthesis, serine palmitoyltransferase (SPT) long chain base subunit 2 (SPTLC2), was the top overall gene hit scoring 6.42 PV. The collagen type IV alpha-3-binding protein (COL4A3BP, 4.92 PV), also known as ceramide transfer protein (CERT) or StAR-related lipid transfer protein 11 (STARD11) was the 10^th^ best hit and the regulator subunit for all three SPTs, serine palmitoyltransferase small subunit A (SPTSSA, 4.47 PV) was the 12^th^ strongest hit. All three of the aforementioned genes, which are key sphingolipid metabolism genes (30), were sensitizing hits – meaning when knocked down they conferred hypersensitivity to the cell fitness-reducing effects of each CAD.

To validate our screen results with SPTLC2 we used myriocin, also known as ISP-1, which is a potent small molecule SPT inhibitor (31) (Figure 1E). We used myriocin and each CAD at doses where they do not affect cell fitness as single agents. Consistent with the sign convention in our screens indicating that SPTLC2 knockdown strongly sensitizes cells to CADs, myriocin had a near complete synthetic lethal interaction with chloroquine and the other CADs we tested (Figure 1F). The level of synthetic interaction we detected between CADs and inhibiting SPT is reminiscent of our results where we identified the well-accepted bisphosphonate mechanism of action (MoA) target farnesyl diphosphate synthase (FDPS) as the top hit when evaluating multiple bisphosphonates (2, 32). These results therefore suggest that inhibiting SPT might be important to the MoA for diverse CADs.

### The overlap in hits from genome-wide screens with pathogens and CADs vs. non-CADs

Considering the top scoring genes with chloroquine were largely indistinguishable from those with other CADs, and chloroquine appears effective against different types of pathogens (namely, parasites and viruses (19, 20)), we wondered which genome-scale screens involving pathogens would overlap best with the CADs. In the last decade there have been numerous genome-scale screens to identify host factors responsible for the toxic effects of pathogens including Ebola (6), adeno-associated AAV2 (7), HIV (8), Rotavirus (9), Influenza A (10), Zika virus (11), Picornavirus (12), Exotoxin A (13), Cholera toxin (14), Type III secretion system and Shiga toxin (15, 16), Ricin toxin (17), and Toxoplasma gondii (18). We analyzed these 14 screens by counting the number of pathogens where each human gene scored as statistically significant. Genes with the same number of pathogen screen hits were further stratified based on their ranking within each screen. This analysis identified sphingolipid metabolic factors including SPTLC2, its cofactor SPTSSA, and the COG and GARP complexes as nine of the top ten overall hits (9/10) (Figure 2B). Moreover, the hits were specific to the CADs and largely didn’t score with non-CADs. The overlap of the top pathogen and CAD screen hits was 80% (24/30) (Figure 2C). Taken together, these results suggest CADs in general would be expected to strongly modulate the toxic effects of diverse pathogens in vitro.

**Figure 2:**
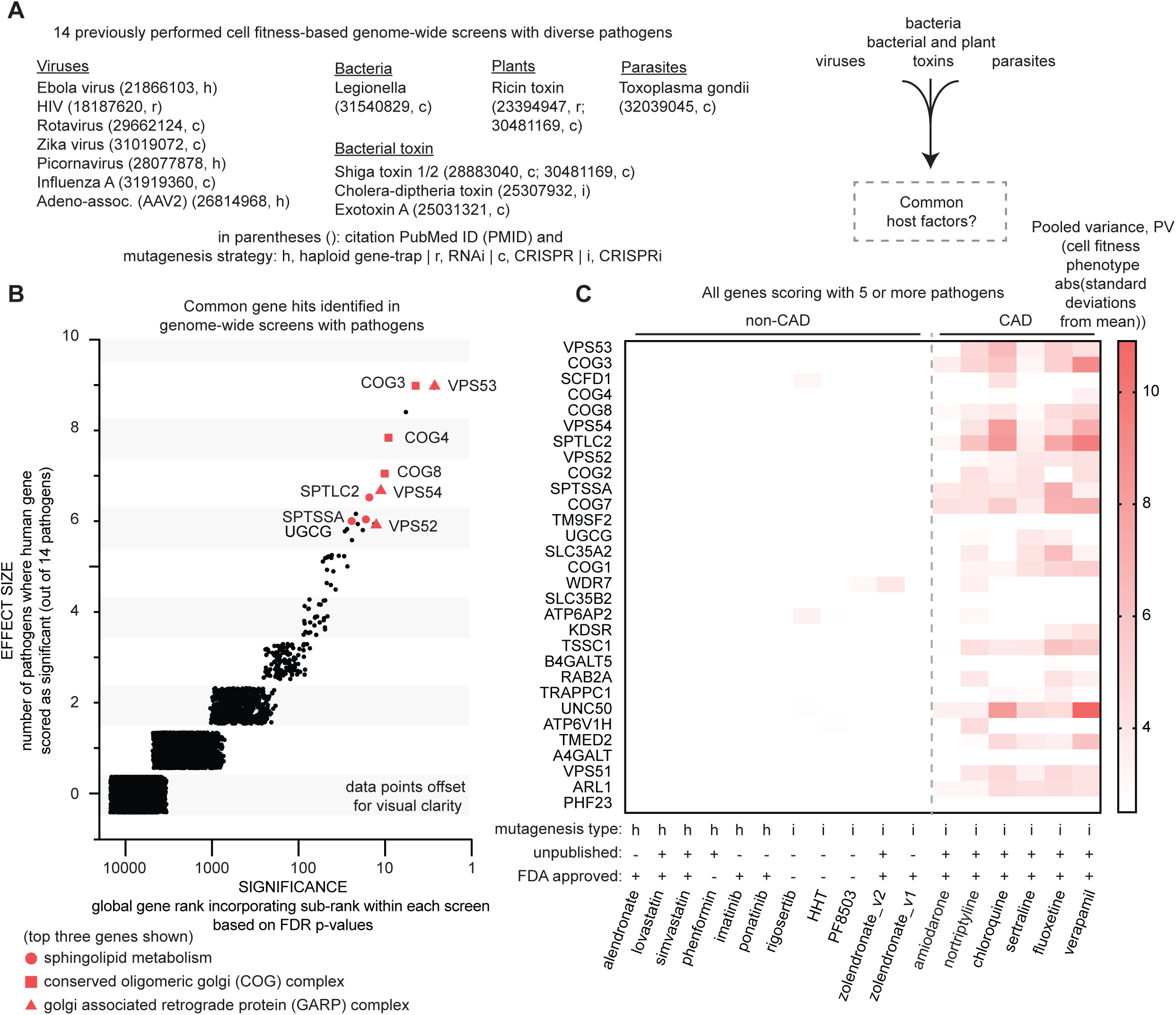
Comparison of genome-wide screens with small molecules with genome-wide screens with pathogens reveals overlap with CADs. **(a)** A list of the pathogens examined in genome-wide screens and their citation identifiers and mutagenesis strategies. **(b)** Visualization of the ranking of 18,903 human genes screened with 14 pathogens. The data points were “jittered” to make them individually visible as there are no genes associated with fractional numbers of pathogens. Genes that scored as significant with the same number of pathogens were further stratified based on their relative positions in each screen. **(c)** Scoring of top pathogen hits in each CAD and non-CAD screen. The top common host genes that scored across pathogens were analyzed as in Figure 1D for their phenotypes with the non-CADs and CADs. The 80% overlap between pathogen and CAD screen hits was determined by assessing how many of the top 30 pathogen screen hits scored with a PV > 2.5 with at least two CADs. As one can visualize from the Figure 2C heatmap, 24 out of 30 genes met this criteria.

### Potential host response and virulence mitigating effects of CADs

Pathogen damaged vesicles get targeted for autophagy to defend cells from the pathogen (33). This is notable because sphingolipid metabolism and the COG and GARP complexes are all related in that they regulate the same step in autophagy, autophagosome formation (34–36). Consistent with this, CADs regulate LC3 cleavage, a key initiating event in autophagy (37), much more strongly than non-CADs (Figure 3A). COGs and GARPs regulate sphingolipid metabolism (38, 39) and viral pathogenicity via autophagy (40). This suggests that one potentially important way CADs may modulate pathogen toxicity is via their effect on host cell autophagy.

**Figure 3:**
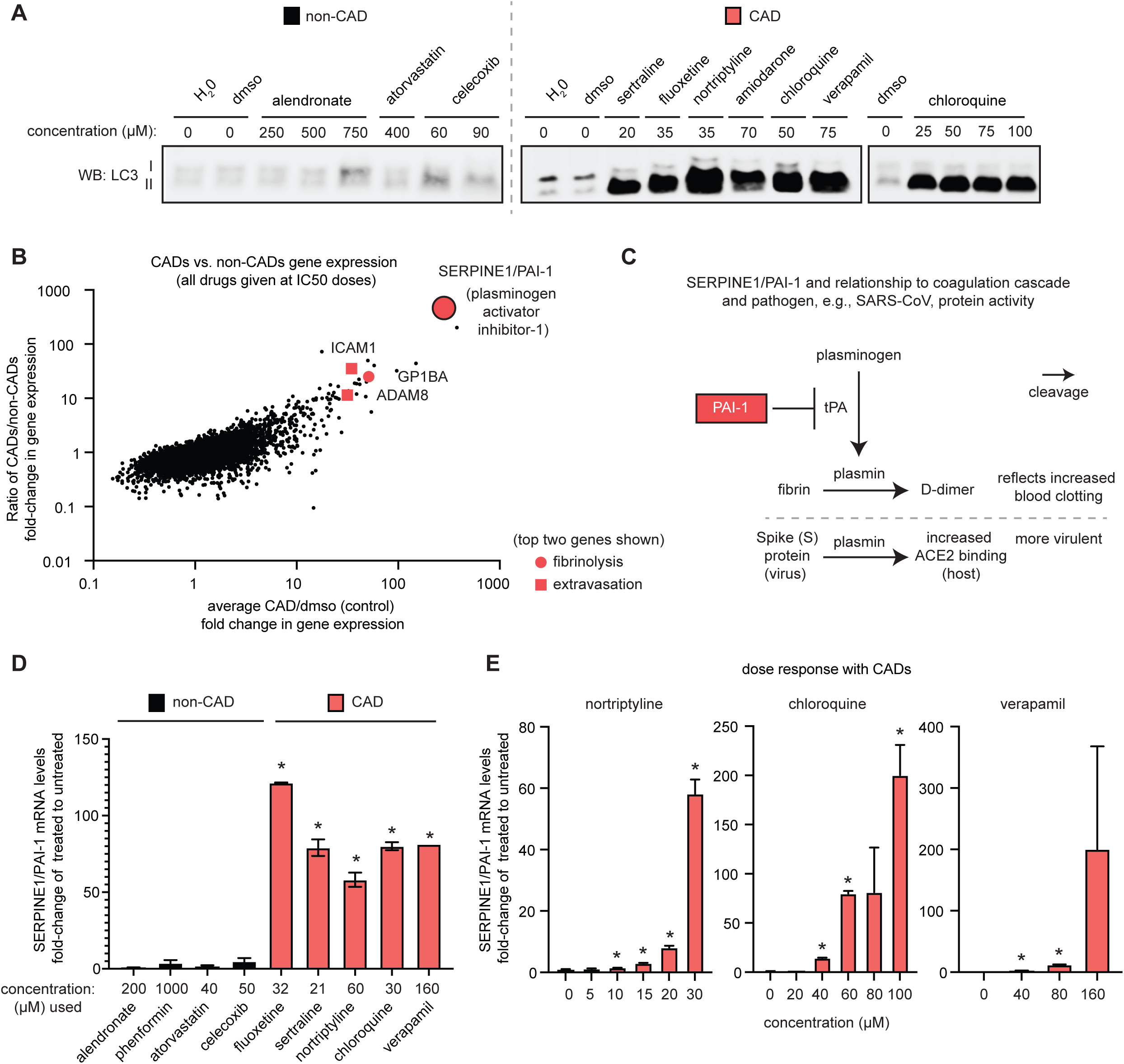
Potential host response and virulence factor-mitigating molecular mechanisms of CADs. **(a)** The effect of CADs and non-CADs on autophagy. K562 cells were treated with the indicated CAD or non-CAD at the indicated concentrations for 24 hours. LC3 cleavage was assessed by immunoblotting. **(b)** Transcriptional profiling of CADs and non-CADs. K562 cells were treated as in (a). The GI50 concentrations of all drugs used were as follows: CADs: sertraline 20 µM, fluoxetine 35 µM, nortriptyline 35 µM, amiodarone 70 µM, chloroquine 50 µM, verapamil 150 µM; non-CADs: phenformin 3000 µM, atorvastatin 400 µM, celecoxib 90 µM, alendronate 250 µM. **(c)** Schematic of the relationship between SERPINE1/PAI-1 and the coagulation cascade and pathogen virulence, and specifically, between SERPINE1/PAI-1-mediated inhibition of plasmin production and D-dimers and pathogen protein cleavage. Plasmin-mediated cleavage of SARS CoV S Protein and its relationship with ACE2 is shown as an example. **(d).** Assessment of SERPINE1/PAI-1 expression in response to CAD or non-CAD treatment. Cells were treated as in (a). SERPINE1/PAI-1 mRNA levels were assessed by RT-qPCR. * P-value < 0.05, one-way ANOVA for all non-CAD vs. CAD samples. Detailed information on statistical tests and exact p-values for each comparison are provided in Table S3. **(e)** Dose dependency of SERPINE1/PAI-1 expression with CADs and non-CADs. Cells were treated as in (a). * P-value < 0.05, unpaired t-test corrected for multiple comparisons by the Holm-Sidak method comparing untreated vs. each CAD dose. Detailed information on statistical tests and exact p-values for each comparison are provided in Table S3.

To gain additional mechanistic insight into the common effects of diverse CADs, we performed transcriptomic analysis of CADs and non-CADs. This identified several non-cell autonomous processes, such as extravasation and fibrinolysis, as top gene ontology categories that were enriched across all six CADs (**Table S3**). Extravasation in the case of immune cells refers to their movement out of the circulatory system and towards the site of tissue damage or infection. Fibrinolysis is a process that prevents blood clots from growing. These gene enrichments seemed interesting to us because like with sphingolipids and the COG and GARP complexes intracellularly, fibrinolysis and extravasation are molecularly linked extracellularly (41).

Among the fibrinolysis hits, SERPINE1/PAI-1 (serine protease inhibitor (serpin) family E member 1/plasminogen activator inhibitor-1) expression showed the largest fold-change vs. control being on average 343-fold increased across the six CADs (Figure 3B). SERPINE1/PAI-1 inhibits tissue plasminogen activator (tPA), which is a key generator of plasmin that drives fibrinolysis (Figure 3C). That our results suggest CADs putatively inhibit fibrinolysis via SERPINE1/PAI-1 seems generally counterintuitive from a therapeutic perspective. Nevertheless, this got our attention because COVID-19 prognosis is associated with elevated D-dimers (42–48), which result from plasmin cleavage (Figure 3C). Plasmin also got our attention because it cleaves the Spike (S) protein in the SARS-CoV-2 related virus, SARS-coV-1, which facilitates its entry via its host receptor angiotensin-converting enzyme 2 (ACE2) (49) – ACE2 is the host receptor for both SARS-coV-1 and SARS-coV-2 (50). (Figure 3C). A potentially broader role of SERPINE1/PAI-1 in pathogenicity is supported by plasmin cleaving other pathogen proteins, such as hemagglutinin of influenza virus, to increase their virulence (51, 52). We validated that PAI-1 expression was specifically elevated by CADs but not non-CADs and determined that the elevation was dose dependent (Figure 3D and E). There is a reciprocal relationship between sphingolipids, SERPINE1/PAI-1, and autophagy. SERPINE1/PAI-1 regulates sphingolipid metabolism (53) and autophagy regulates SERPINE1/PAI-1 (54). Therefore, potentially related to regulating host responses such as autophagy, these results suggest CADs and sphingolipids might influence pathogen virulence via SERPINE1/PAI-1.

## DISCUSSION

These results are important in light of novel coronavirus SARS-CoV-2 and COVID-19 because each CAD we tested is already FDA approved. One of the CADs we screened, amiodarone, has shown efficacy in vitro against SARS-CoV-1 (55). Another CAD we screened, verapamil, was recently identified by Gordon et al. using proteomics analysis as possibly being relevant to SARS-CoV-2 (56) (not yet peer reviewed). This suggests amongst the drugs Gordon et al. discovered, verapamil might potentially be prioritized for further pre-clinical investigation. Lastly, azithromycin has shown promise in combination with chloroquine against SARS-CoV-2 (57). This is interesting considering azithromycin’s pKa (8.5) and cLogP (2.6) and ability to promote phospholipidosis in vivo (58), which suggests it might also be acting as a CAD in this context. Some COVID-19 patients are responding well to chloroquine whereas others are not. As the pandemic continues it will be important to better understand who responds to treatment. We established that genome-wide screening in cells can identify key genetic determinants that predict bad outcomes from taking bisphosphonates (1). A similar cellular analysis with SARS-CoV-2 treatments should be considered as soon as there are clear frontrunners.

In addition to SARS-CoV-2, our work suggests that CADs could be considered for further pre-clinical investigation for other pathogens including those yet-to-emerge which do not have an effective therapy or vaccine. Indeed, many CADs including those we have not studied here, such as tamoxifen, are being increasingly noted for their inhibitory effects on various infectious agents even in vivo (59–61).

To our knowledge, this is the first meta-analysis of genome-wide screens in human cells to identify consensus host factors important to human pathogens. Most genome-wide screens in human cells with pathogens have focused on poorly characterized genes rather than potentially universally important ones. Identifying “universal” host factors such as sphingolipid metabolism and the COG and GARP complexes is important in light of the recently put forth Omnigenic model (62). This model states that most genes contribute to human disease, though “core” genes contribute the most. There isn’t a clear understanding of what might be a core gene for most diseases. Our work suggests sphingolipid metabolism and the COG and GARP complexes represent potential core host response genes for diverse human pathogens.

There are several potential reasons related to membrane biology that may explain why CADs and sphingolipids are particularly important in host responses to pathogens: 1) Sphingolipids are a key part of lipid rafts (63), which are a common point of entry for many pathogens (64). For example, SARS-CoV-1 requires sphingolipids in lipid rafts for binding to ACE2 (65); 2) Once inside the cell, pathogens are also susceptible to the unique chemical properties of sphingolipids and CADs. For example, ceramides are non-swelling amphiphiles implying that they cannot give rise to micelles or other aggregates (66). Though understudied, this would seem to modulate the membranes of inclusion bodies, which are used by many pathogens to evade their destruction (67). Also, some pathogens need the ER to make more of themselves (68). Considering CADs associate with membranes (69) potentially in the ER, this could explain why CADs are effective against these types of pathogens; 3) CADs potentiate the effects of many anti-pathogen drugs. For example, zinc is an anti-viral when used in combination therapies (70). The internalization of zinc ions is largely limited by their incompatibility with the cell membrane chemistry. Chloroquine is a zinc ionophore which increases zinc’s uptake by cells thereby promoting its biological effects (71).

There are several limitations from our analysis: 1) Like chloroquine, myriocin is an immunosuppressant (72). This suggests if myriocin eventually becomes considered as a treatment against pathogens, it might make a patient’s situation worse. Consistent with this, in preclinical studies myriocin has been shown to promote host infection by herpes simplex (73) and dengue viruses (74); 2) We only compared genome-wide mutagenesis screens where cell fitness had been used as the phenotype. We acknowledge reporter-based screens can yield important information we might be missing. However, we used cell fitness because it is robust, and more importantly, because it enabled us to make apples-to-apples comparisons between screens. This is critical for comparing screens using different mutagenesis strategies (CRISPR, CRISPRi, haploid, RNAi) and different types of perturbations (small molecules and pathogens) as we’ve done here; 3) Our results suggest CADs inhibit SPT thereby inhibiting sphingolipid biosynthesis and ceramide production. Previous studies with antidepressants demonstrated that CADs act on sphingolipid catabolism, in particular on acid sphingomyelinase (24), which also generates ceramide. We cannot rule out that CADs decrease ceramide levels both at the level of anabolic and catabolism processes. However, our screens strongly suggest the former.

Prions are a type of pathogen we did not include in our analysis because a genome-wide screen in human cells has not yet been done with one. Considering prions or other misfolded pathogenic proteins such as amyloid and tau also engage autophagy (75), we hypothesize sphingolipid biosynthesis and the COG and GARP complexes would also score strongly in a genome-wide screen with such proteins. There is growing evidence that infectious agents might be an important causative factor in Alzheimer’s disease (AD) (76). Infectious agents and misfolded proteins are both by definition, pathogens. In the future, it will be interesting to assess CADs for their effects on the interrelations of both types of pathogens in the development of AD or other neurodegenerative conditions.

## Supporting information

CADs_pathogens_Supp. Table 1

CADs_pathogens_Supp. Table 2

CADs_pathogens_Supp. Table 3

## Abbreviations

CADs: cationic amphipathic (amphiphilic) drugs, also known as lysosomotropic agents
SARS-CoV-1: Severe acute respiratory syndrome coronavirus-1
SARS-CoV-2: Severe acute respiratory syndrome coronavirus-2
COVID-19: Coronavirus Disease-2019 where SARS-CoV-2 is the causative agent
FDA: US Food and Drug Administration
MOA: mechanism of action
SPT: serine palmitoyltransferase
SPTLC2: serine palmitoyltransferase long chain base subunit 2
COG: conserved oligomeric golgi complex
GARP: golgi-associated retrograde protein complex
SERPINE1/PAI-1: serine protease inhibitor (SERPIN) Family E Member 1/plasminogen activator inhibitor-1

---

**Figure S1:**
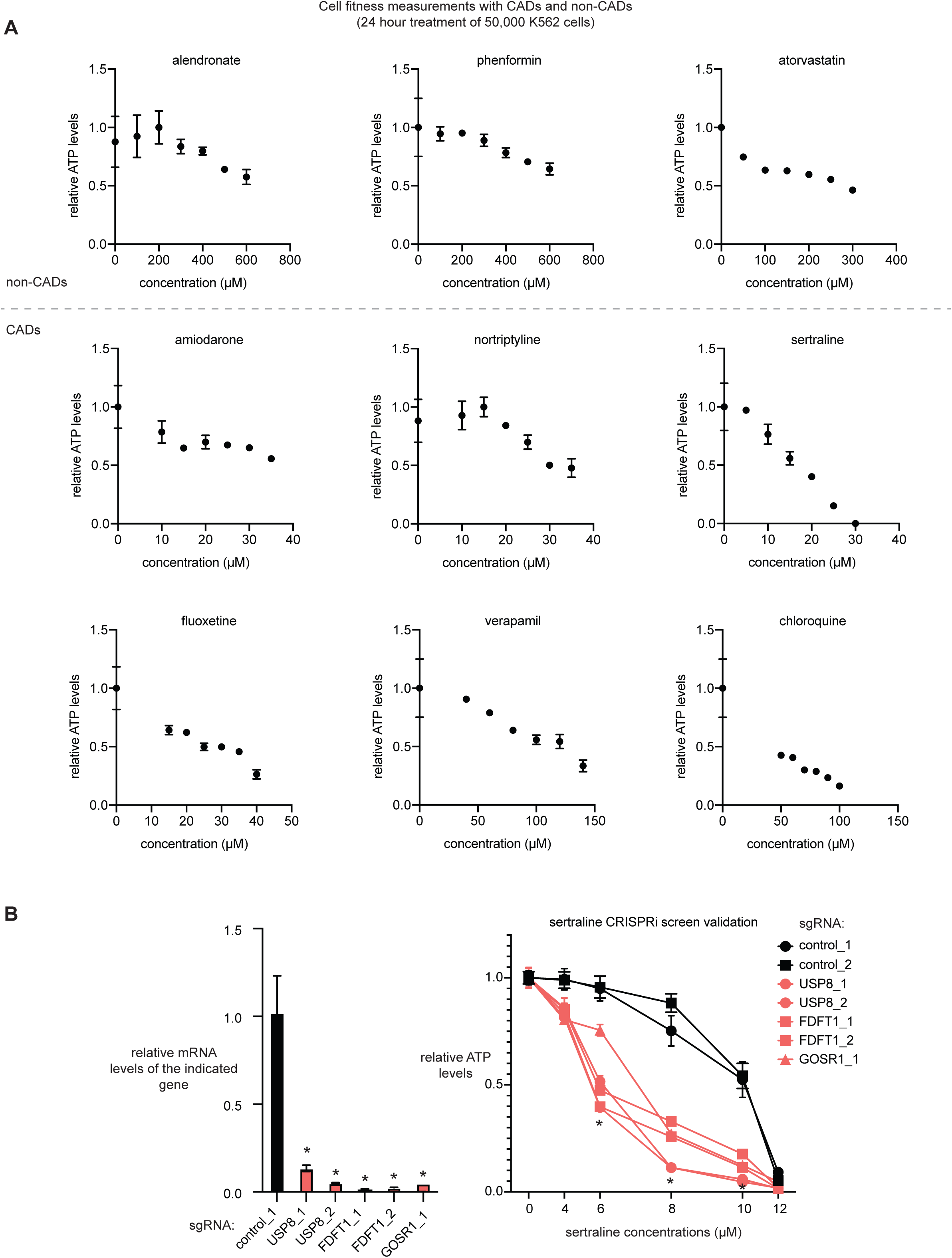
Data related to Figure 1. **(a)** Growth inhibition dose response curves for CADs and non-CADs. 50,000 K562 cells were treated with the indicated drugs for 24 hours. ATP levels were used as a proxy for cell fitness. **(b)** Example validation of gene hits from the sertraline CRISPRi screen. Relative mRNA and ATP levels for control and knockdown for the genes USP8, FDFT1, and GOSR1 in K562 cells in the presence of the indicated concentrations of sertraline or vehicle control DMSO are shown. ATP levels were analyzed following 72 hour treatment. mRNA levels were assessed by RT-qPCR. * P-value < 0.05, ANOVA for all controls vs. experimental samples. One-way or two-way ANOVA was performed for mRNA expression or ATP analysis, respectively. Detailed information on statistical tests and exact p-values for each comparison are provided in Table S1.

**Table S1: Data related to Figure 1.** The following tables related to Figure 1 and Figure S1 are included in Table S1: All genome-wide screen results from this work; Further chemical property information for CADs and non-CADs; Details on previously published genome-wide small molecules screens that were included in our analysis; Statistical analysis performed in Figure 1 and S1 including exact p-values.

**Table S2: Data related to Figure 2.** The following tables related to Figure 2 are included in Table S2: Details on each pathogen genome-wide screens analyzed. Enrichr-determined Gene ontology categories for the genes identified as most commonly scoring across the 14 pathogen screens.

**Table S3: Data related to Figure 3.** The following tables related to Figure 3 are included in Table S3: Full RNA-Seq data with CADs and non-CADs. Enrichr-determined gene ontology provided for genes 5X or more enriched with all CADs. Statistical analysis performed in Figure 3 including exact p-values.

## ACKNOWLEDGMENTS

We thank current and past members of the Peterson for helpful discussions and general assistance, especially S. Diemar, C. Chow, and K. Myachin. We thank E. Gulbins, T. Hornemann, G. D’Angelo, J. Shayman for discussions on sphingolipids.

## AUTHOR CONTRIBUTIONS

T.R.P. and S.K. designed the study. S.K., J.L., and N.J.S. performed the CRISPRi screens with CADs. T.R.P., J.C., M.V., and T.R.B. performed the haploid screens with non-CADs. S.K., J.P., D.T.B., T.H.P., and I.C.B. performed screen validation. S.K. and N.B. performed the myriocin experiments. N.C.J. and A.C-O. analyzed PubMed to investigate existing evidence of the usage of CADs with pathogens. A.C-O. performed statistical analysis to obtain the GI50s for all drugs. T.R.P. performed the pathogen meta-analysis. S.K. and S.K.H. performed the autophagy experiments. S.K. performed the RNA-Seq and SERPINE1/PAI-1 experiments. R.D. provided chemical analysis on the CADs and non-CADs. T.R.P. wrote the paper.

## DATA AND MATERIALS AVAILABILITY

All data associated with this study are present in the paper or the Supplementary Materials. Shared reagents are subject to a materials transfer agreement.

## MATERIALS & METHODS

### Materials

Reagents were purchased from the following manufacturers: DMEM and RPMI 1640 Medium, GlutaMAX Supplement from GE Healthcare Gibco; Fetal Bovine Serum (FBS) and from ThermoFisher Scientific HyClone; Transit LT-1 reagent (cat. # MIR 2300) from Mirus Bio; Polyethylenimine linear MW 25000 transfection grade (PEI 25K) (cat. # 239661) from Polysciences Inc.; Chloroquine diphosphate (Cat. # C6628), sertraline (cat. # S6319), verapamil (cat. # V4629), fluoxetine (Cat. # 1279804), amiodarone (cat. # PHR1164), nortriptyline (cat. # N7261), metformin (cat. # PHR1084), phenformin (cat. # PHR1573), alendronic acid (cat. # 1012780), atorvastatin (cat. # PHR1422), lithium carbonate (cat. # 203629) from Millipore Sigma; Bolt 4-12% Bis-Tris gels, Halt Protease Inhibitor Cocktail from Invitrogen; Bradford Reagent from Bio-Rad; LC3B antibody (cat. # 2775S) from Cell Signaling Technology; Chamelon Duo Pre-Stained Ladder, IRDye 800CW and IRDye 680RD secondary antibody from Li-COR; Cell-titer Glo (cat. # G7572) from Promega; NEBNext Ultra II Q5 Master Mix (cat. # M0544L), BstXI (cat. # R0113L), Blp I (cat. # R0585L) from New England Biolabs; NucleoSpin Blood DNA isolation kit (cat. # 740950.50) from Macherey Nagel; Human Genome-wide CRISPRi-v2 Libraries were provided by Jonathan Weissman via Addgene (cat. # 1000000090).

### Cell Lines and Tissue Culture

K562 human myeloid leukemia cells and human embryo kidney (HEK) 293T cells were obtained from ATCC. HEK 293T cell lines were cultured in DMEM with 10% FBS and 1% penicillin/streptomycin. K562 cell lines were cultured in RPMI1640 Medium GlutaMAX Supplement with 10% FBS and 1% penicillin/streptomycin. All cell lines were maintained at 37°C and 5% CO_2_.

### Genome-scale screens

Genome-scale screens were carried out similar to the previously published screens (2, 3, 14, 78, 79). CRISPRi v2 sgRNA libraries (78) were transduced into K562 CRISPRi-competent (dCas9-KRAB) cells at a low MOI (~0.3). Two days after transduction, the infected cells were selected with 0.75 ug/ml puromycin for three days, and the transduction was confirmed by flow cytometry. Cells were recovered from puromycin selection for two days. After two days of recovery, initial sample cells (T0; 200 million) were frozen down, and remaining cells (400 million cells) were split into either untreated (C) or treated (D) with the drug of interest. For each flask in the untreated and treated groups, the cells were kept to 0.5 million cells/ml daily. Drug treatment was continued until the untreated cells doubled five to eight more times than the treated cells. Cells were then recovered for 1 week to allow the treated cells to undergo three to four doublings. Genomic DNA isolation and library preparations were performed as previously described (14).

Genome-scale gene trap screens using haploid KBM7 cells were performed as previously described (77). In brief, the genetic selection with the indicated small molecule drug was performed on 100 million mutagenized KBM7 cells (80). Cells were exposed to drug and allowed to recover for several weeks (on average, 4 weeks) before harvesting the surviving cells and isolating genomic DNA. Analysis of the screen is as follows: the sequences flanking retroviral insertion sites were determined using inverse PCR followed by Illumina sequencing. After mapping these sequences to the human genome, we counted the number of inactivating mutations (mutations in the sense orientation or present in exon) per individual Refseq-annotated gene as well as the total number of inactivating insertions for all Refseq-annotated genes. For each gene, statistical enrichment was calculated by comparing how often that gene was mutated in the drug-treated population with how often the gene carries an insertion in the untreated control dataset. For each gene, the *P*-value (corrected for false discovery rate) was calculated using the one-sided Fisher exact test.

#### sgRNA manipulations

sgRNA cloning for CRISPRi screen validation studies was performed according to the Weissman lab protocol: https://weissmanlab.ucsf.edu/links/sgRNACloningProtocol.pdf.

For individual validation of sgRNA phenotypes, sgRNA protospacers targeting the indicated genes or control protospacers target eGFP or non-targeting, control_1 and control_2 (we colloquially refer to these controls as “PBA392” and “PMJ051”, respectively), were individually cloned by annealing complementary synthetic oligonucleotide pairs (Integrated DNA Technologies) with flanking BstXI and BlpI restriction sites and ligating the resulting double-stranded segment into BstXI/BlpI-digested pCRISPRia-v2 (marked with a puromycin resistance cassette and BFP, Addgene #84832; (78)). Protospacer sequences used for individual evaluation are listed below. The resulting sgRNA expression vectors were individually packaged into lentivirus. Internally controlled growth assays to evaluate sgRNA drug sensitivity phenotypes were performed by transducing cells with sgRNA expression constructs at MOI < 1 (15 – 30% infected cells), selecting to purity with puromycin (0.75 μg/mL), allowing to recover for at least 1 day, treating cells with the indicated concentrations of drugs or DMSO 4-7 days after infection, and measuring the fraction of sgRNA-expressing cells 72 hours after that. During this process, populations of cells were harvested for measurement of mRNA levels by RT-qPCR (see below). These experiments were performed in triplicates from the treatment step. Knockdown of each gene was performed with their own batch of control sgRNA cells, and the data from the control cells were averaged to allow comparison of the genes on the same scale.

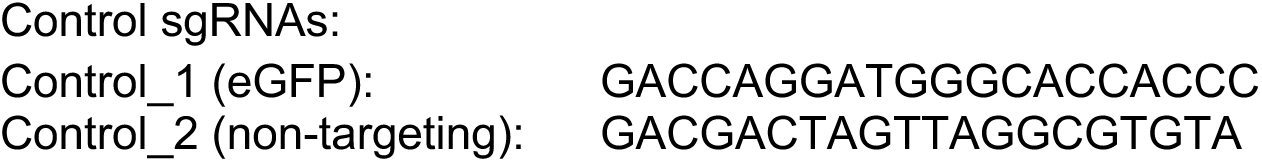

CRISPRi sgRNAs sequences below were obtained from the aforementioned CRISPRi-v2 library. We selected two out of the 10 available sgRNAs based on which ones scored best in screening.

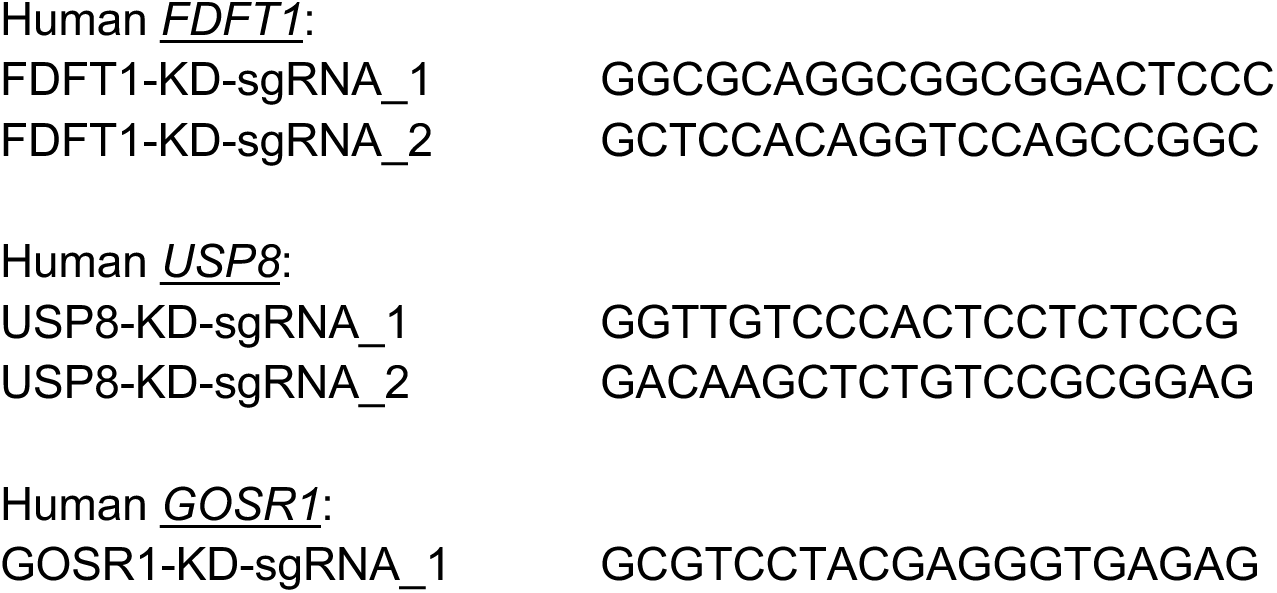

#### Gene expression analysis

For individual gene analysis, total RNA was isolated and reverse-transcription was performed from cells or tissues in the indicated conditions. The resulting cDNA was diluted in Dnase-free water (1:20) followed by quantification by real-time PCR. mRNA transcripts were measured using Applied Biosystems 7900HT Sequence Detection System v2.3 software. All data was expressed as the ratio between the expression of target gene to the housekeeping genes ACTB (actin) and/or GAPDH. Each treated sample was normalized to controls in the same cell type.

Human PCR primer sequences were obtained from: https://pga.mgh.harvard.edu/primerbank/.

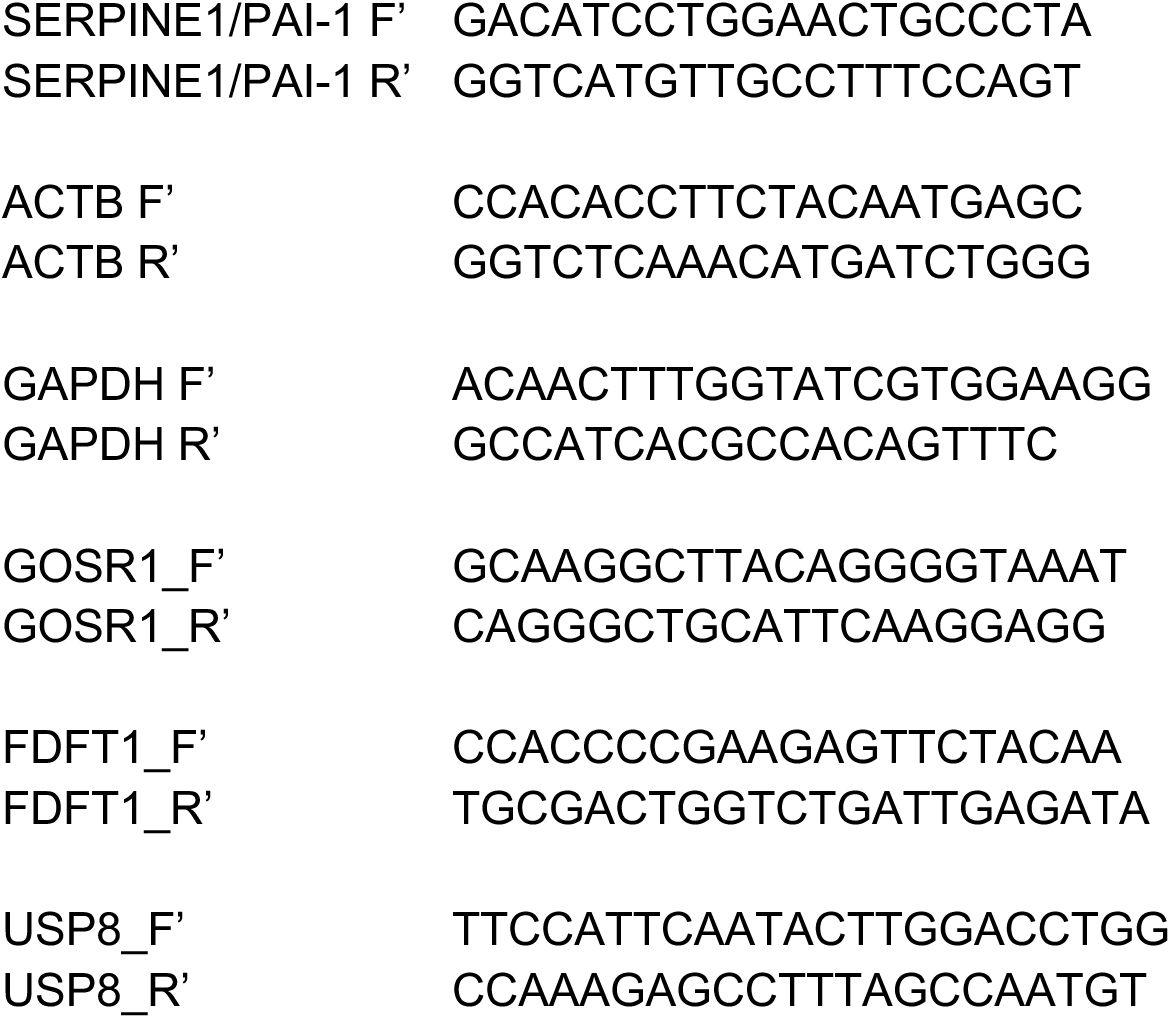

For RNA-Seq experiments, total RNA was prepared from K562 cells and analyzed using standard pipelines.

#### Cell fitness assays

Wild-type or mutant cells were seeded at 10,000 or 50,000 cells per well in a 96-well tissue culture plate and treated with indicated concentrations of compound or left untreated. 24 or 72 hours after treatment the cell viability was measured using a Cell-titer Glo colorimetric assay (Promega) according to manufacturer’s protocol. Fitness was plotted as percentage compared to untreated control. Growth Inhibitory 50% GI50 and not Inhibitory Concentration 50% IC50 was used because the latter refers to 50% inhibition of the maximal inhibition. The maximum inhibition varies for each drug, therefore using GI50 instead allowed us to compare all drugs on the same scale.

#### Protein analysis

All cells were rinsed twice with ice-cold PBS and lysed with cell lysis buffer (Cell Signaling) in which Halt™ Protease and Phosphatase Inhibitor Cocktail (#78440 ThermoFisher Scientific) was added. Lysate was incubated at 4 centigrade for 15-30min with constant inversion. The soluble fractions of cell lysates were isolated by centrifugation at 13,000 rpm for 10 min in a microcentrifuge. Lysate protein concentrations were normalized by Bradford assay (Bio-Rad). Proteins were then denatured by the addition of sample buffer and by boiling for 5 minutes, resolved using 4%-20% SDS-PAGE (Invitrogen), and analyzed by immunoblotting for the indicated proteins. Immunoblotting was performed as follows: nitrocellulose membranes were blocked at room temperature (RT) with 5% BSA for 45min. Membranes were then incubated overnight at 4°C with indicated primary antibodies dissolved in 5% BSA. Membranes were then washed 3 times in TBST, each wash lasting 5min. Membranes were then incubated at RT with desired secondary antibodies at 1:2000 in 5% milk for 45min. HRP-conjugated or fluorescent secondary antibodies (Santa Cruz Biotechnology or Thermo Fisher, respectively) were used for detection. Membranes were then washed 3 times in TBST, each wash lasting 5min. Signal from membranes using HRP-conjugated secondary antibodies were captured using a camera and those using fluorescent secondary antibodies were imaged on a GE Typhoon Trio imager.

### Cheminformatic & bioinformatic analysis

Chemical properties for CADs and non-CADs were obtained from PubChem: https://pubchem.ncbi.nlm.nih.gov/ or DrugBank: https://www.drugbank.ca/ when PubChem was incomplete. The additional chemical properties presented in Table S1 were calculated using Advanced Chemistry Development (ACD/Labs) Software V11.02. Images of chemical structures were obtained from Wikipedia.

The chemical differences between the non-CADs (bisphosphonates, biguanides, and statins) used throughout this work as a follows: Bisphosphonates – alendronate differs from zoledronic acid (zoledronate) with its R^2^ group. Biguanides – phenformin, which was used throughout the work, differs from metformin in having a phenyl ring. Unlike in vivo where both phenformin and metformin regulate metabolism consistent with the expected MoA for biguanides, in vitro only phenformin is active at putatively relevant concentrations (µM). The reasons for the poor metformin efficacy in vitro aren’t well understood. Statins – The statins used in this work: atorvastatin, lovastatin, and simvastatin are all relatively lipophilic compared to the hydrophilic statins such as rosuvastatin. Simvastatin is more closely structurally related to lovastatin than to atorvastatin.

Amongst all previously published small molecule and pathogen screens, only gene loss of function (i.e., haploid gene-trap, CRISPR, RNAi) and not gain-of-function screens (i.e., CRISPRa, retroviral overexpression) were considered in our analysis. Multiple mutagenesis strategies were compared to ensure any genes considered hits weren’t only found with one method. Screens without readily apparent means to separate significant gene hits vs. non-significant genes were not considered. Our rationale for choosing the non-CADs we analyzed are as follows: rigosertib, HHT, and PF8503 are, to our knowledge, the only non-CADs that have been analyzed using CRISPRi in K562 cells, which is the mutagenesis strategy and cell type we used with CADs. Ponatinib and imatinib are non-CADs that were screened in HAP1 cells, which is one of the common cell types that we analyzed (see Table S1) because it has been commonly used by the research community to identify host factors for pathogens. Examples of screens for both pathogens and small molecule screens that we did not analyze include:

- Ebola (pathogen) – PMID: 30655525 – unable to understand the reported statistical significance cut-off.
- T. cruzi (pathogen) – PMID: 21625474. This was a visual-based screen, whereas we focused on cell fitness-based screens.
- Sorafenib (small molecule) – PMID: 29467456 – unable to identify a statistical significance cut-off.

Gene ontology analysis was performed using Enrichr: https://amp.pharm.mssm.edu/Enrichr/. Genes of interest were input into Enrichr and output was taken from “Ontologies” -> ”Go Biological Process 2018”. Similar results were obtained using the Gene Ontology web service: http://geneontology.org/, though that service’s algorithm prioritizes larger clusters of genes in calculating significant hits.

For Figure 1 CRISPRi analysis, there were 19 genes input to Enrichr that were increased by 2.5x with treatment relative to DMSO vehicle control. For Figure 3 gene expression analysis, there were 48 genes input to Enrichr that were increased by 5x with treatment relative to DMSO vehicle control where the DMSO counts per million (CPM) was greater than 1.

### Statistical and graphical analysis

Unless otherwise specified, groups were compared by analysis of variance (ANOVA) correcting for multiple comparisons with the Tukey method using GraphPad Prism version 8.4.0 software. All data are expressed as the mean ± s.d. GraphPad Prism was also used to generate all graphs except figure 2B where RStudio and ggplot2 was used. All code is available via the git repository: https://github.com/tim-peterson/CADs-pathogens (DOI: 10.5281/zenodo.3742662). To combine p-values for individual screens as done in Figure 1C, a false discovery rate (FDR, also known as Benjamini & Hochberg method) adjusted p-value was used. The python code below was used. Imported libraries are omitted for clarity, but the code that invokes them is available via the ‘CADs-pathogens’ git repository:

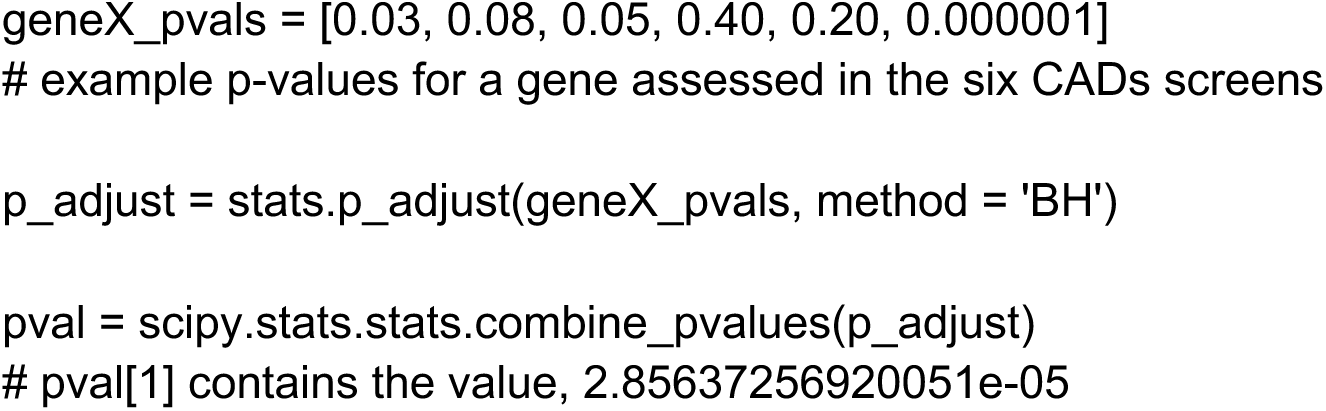

